# Language Models Outperform Cloze Predictability in a Cognitive Model of Reading

**DOI:** 10.1101/2024.04.29.591593

**Authors:** Adrielli Lopes Rego, Joshua Snell, Martijn Meeter

## Abstract

Although word predictability is commonly considered an important factor in reading, sophisticated accounts of predictability in theories of reading are yet lacking. Computational models of reading traditionally use cloze norming as a proxy of word predictability, but what cloze norms precisely capture remains unclear. This study investigates whether large language models (LLMs) can fill this gap. Contextual predictions are implemented via a novel parallel-graded mechanism, where all predicted words at a given position are pre-activated as a function of contextual certainty, which varies dynamically as text processing unfolds. Through reading simulations with OB1-reader, a cognitive model of word recognition and eye-movement control in reading, we compare the model’s fit to eye-movement data when using predictability values derived from a cloze task against those derived from LLMs (GPT2 and LLaMA). Root Mean Square Error between simulated and human eye movements indicates that LLM predictability provides a better fit than Cloze. This is the first study to use LLMs to augment a cognitive model of reading with higher-order language processing while proposing a mechanism on the interplay between word predictability and eye movements.

**Author Summary:** Reading comprehension is a crucial skill that is highly predictive of later success in education. One aspect of efficient reading is our ability to predict what is coming next in the text based on the current context. Although we know predictions take place during reading, the mechanism through which contextual facilitation affects ocolarmotor behaviour in reading is not yet well-understood. Here, we model this mechanism and test different measures of predictability (computational vs. empirical) by simulating eye movements with a cognitive model of reading. Our results suggest that, when implemented with our novel mechanism, a computational measure of predictability provide better fits to eye movements in reading than a traditional empirical measure. With this model, we scrutinize how predictions about upcoming input affects eye movements in reading, and how computational approches to measuring predictability may support theory testing. In the short term, modelling aspects of reading comprehension helps reconnect theory building and experimentation in reading research. In the longer term, more understanding of reading comprehension may help improve reading pedagogies, diagnoses and treatments.

## Main

Humans can read remarkably efficiently. What underlies efficient reading has been subject of considerable interest in psycholinguistic research. A prominent hypothesis is that we can generally keep up with the rapid pace of language input because language processing is predictive, i.e., as reading unfolds, the reader anticipates some information about the upcoming input (1–3). Despite general agreement that this is the case, it remains unclear how to best operationalize contextual predictions (3,4). In current models of reading (5–7)), the influence of prior context on word recognition is operationalized using Cloze norming, which is the proportion of participants that complete a textual sequence by answering a given word. However, Cloze norming has both theoretical and practical limitations, which are outlined below (4,8,9). To address these concerns, in the present work we explore the use of Large Language Models (LLMs) as an alternative means to account for contextual predictions in computational models of reading. In the remainder of this section, we discuss the limitations of the current implementation of contextual predictions in models of reading, which includes the use of Cloze norming, as well as the potential benefits of LLM outputs as a proxy of word predictability. We also offer a novel parsimonious account of how these predictions gradually unfold during text processing.

Computational models of reading are formalized theories about the cognitive mechanisms that may take place during reading. The most prominent type of model are models of eye-movement control in text reading (see (10) for a detailed overview). These attempt to explain how the brain guides the eyes, by combining perceptual, oculomotor and linguistic processes. Despite the success of these models in simulating some word-level effects on reading behaviour, the implementation of contextual influences on the recognition of incoming linguistic input is yet largely simplified. Word predictability affects lexical access of the upcoming word by modulating either its recognition threshold (e.g. EZ-reader (5) and OB1-reader(6)) or its activation (e.g. SWIFT(7)). But both approaches rely on a pre-determined, fixed value for each word, as such not considering the possibility that predictability varies dynamically as text processing unfolds.

What is more, the pre-determined, fixed value in such models is conventionally operationalized with Cloze norming (11). Cloze predictability is obtained by having participants write continuations of an incomplete sequence, and then taking the proportion of participants that have answered a given word as the Cloze probability of that word. The assumption is that the participants draw on their individual lexical probability distributions to fill in the blank (e.g. *house* may be more probable than *place* to complete *I met him at my* for participant A, but not for participant B), and that Cloze reflects some overall subjective probability distribution. However, scientists have questioned this assumption (4,8,9). The Cloze Task is an offline and untimed task, leaving ample room for participants to consciously reflect on sequence completion and adopt strategic decisions (4). This may be quite different from normal reading where only ∼200ms is spent on each word (12). Another issue is that Cloze cannot provide estimates for low-probability continuations, in contrast with behavioural evidence showing predictability effects of words that never appear among Cloze responses, based on other estimators, such as part-of-speech (8,13). Thus, Cloze completions likely do not perfectly match the rapid predictions that are made online as reading unfolds.

Predictability values generated by LLMs may be a suitable methodological alternative to Cloze completion probabilities. LLMs are computational models whose task is to assign probabilities to sequences of words (14). Such models are traditionally trained to accurately predict a token given its contextual sequence, similarly to a Cloze Task. An important difference, however, is that whereas Cloze probability is an average across participants, probabilities derived from LLMs are relative to every other word in the model’s vocabulary. This allows LLMs to better capture the probability of words that rarely or never appear among Cloze responses, potentially revealing variation in the lower range (15). In addition, LLMs may offer a better proxy of semantic and syntactic contextual effects, as they computationally define predictability and how it is learned from experience. The model learns lexical knowledge from the textual data, which can be seen as analogous to the language experience of humans. The meaning of words is determined by the contexts in which they appear (*distributional hypothesis* (16)) and the consolidated knowledge is used to predict the next lexical item in a sequence (9). Finally, language models have recently been shown to perform as well as (17), or outperform(18), predictability estimates derived from Cloze Tasks in fitting oculomotor data. This evidence has led to believe that LLMs may be suitable for theory development in models of eye-movement control in reading (9).

The present study marks an important step in exploring the potential of language models in advancing understanding of the reading brain (19,20), and more specifically, of LLMs’ ability to account for contextual predictions in models of eye-movement control in reading (9). We investigate whether a model of eye-movement control in reading can more accurately simulate reading behaviour using predictability derived from transformer-based LLMs or from Cloze. Assuming that predictability during reading is lexical, non-discrete, parafoveal, parallel and dynamic (see *Reading Simulations* section of Methods for explanation), we hypothesize that LLM-derived probabilities capture semantic and syntactic integration of the previous context, which in turn affects processing of upcoming bottom-up input. This effect is expected to be captured in the early reading measures (see Methods).

Since predictability may also reflect semantic and syntactic integration of the predicted word with the previous context (15), late measures are also evaluated. Importantly however, employing LLM-generated predictions is only one part of the story. A cognitive theory of reading also has to make clear how those predictions operate precisely: i.e., when, where, how and why do predictions affect processing of upcoming text? The aforementioned models have been agnostic about this. Aiming to fill this gap, our answer, as implemented in the updated OB1-reader model, is as follows.

We propose that making predictions about upcoming words affects their recognition through parallel and graded pre-cycle activation. Predictability is graded because it modulates pre-cycle activation of all words predicted to be at a given position in the parafovea to the extent of each word’s likelihood. This means that higher predictability leads to a stronger pre-cycle activation of all words predicted to be at a given position in the parafovea.

Predictability is also parallel, because predictions can be made about multiple text positions simultaneously. With each processing cycle, this predictability-derived activation is summed to the activity resulting from visual processing of the previous cycle and weighted by the predictability of the previous word, which in turn reflects the prediction certainty up to the current cycle (see Methods for more detailed explanation). In this way, predictability gradually and dynamically affects words in parallel, including non-text words in the model’s lexicon.

With these mechanisms in place, we hypothesize that OB1-reader achieves a better fit to human oculomotor data with LLM-derived predictions than with Cloze-derived predictions. To test this hypothesis, we ran reading simulations with OB1-reader either using LLM-derived predictability or Cloze-derived predictability to activate words in the model’s lexicon prior to fixation. The resulting reading measures were compared with measures derived from eye-tracking data to evaluate the model’s fit to human data. This is the first study to combine a language model with a computational cognitive model of eye-movement control in reading to test whether the output of LLMs is a suitable proxy for word predictability in such models.

## Results

For the reading simulations, we used OB1-reader (6). In each processing cycle from OB1-reader’s reading simulation, the predictability values were used to activate the predicted words in the upcoming position (see *Reading Simulations* section in Methods for a detailed description). We ran 100 simulations per condition in a “3×3 + 1” design: three predictability estimators (Cloze, GPT2 and LLaMA), three predictability weights (low = 0.05, medium = 0.1, and high = 0.2) and a baseline (no predictability). For the analysis, we considered eye-movement measures at word-level. The early eye-movement measures of interest were skipping (SK), i.e. the proportion of participants who skipped the word on first pass; first fixation duration (FFD), i.e. the duration of the first fixation on the word; and gaze duration (GD), i.e. the sum of fixations on the word before the eyes move forward. The late eye-movement measures of interest were total reading time (TRT), i.e. the sum of fixation durations on the word; and regression (RG), i.e. the proportion of participants who fixated the word after the eyes have already passed the text region the word is located. To evaluate the model simulations, we used the reading time data from the Provo corpus (21) and computed the Root Mean Squared Error (RMSE) between each eye-movement measure from each simulation by OB1-reader and each eye-movement measure from the Provo corpus averaged over participants.

### Fit across eye-movement measures

In line with our hypotheses, OB1-reader simulations were closer to the human eye-movements with LLM predictability than with Cloze predictability. Fig. 1 shows the standardized RMSE of each condition averaged over eye movement measures and predictability weights. To attest the predictability implementation proposed in OB1-reader, we compared the RMSE scores between predictability conditions and baseline. All predictability conditions reduced the error relative to the baseline, which confirms the favourable effect of word predictability on fitting word-level eye movement measures in OB1-reader. When comparing the RMSE scores across predictability conditions, the larger language model LLaMA yielded the least error. When comparing the error among predictability weights (see Fig. 3f), LLaMA yielded the least error in all weight conditions, while GPT2 produced less error than Cloze only with the low predictability weight. These results suggest that language models, especially with higher prediction accuracy, are good word predictability estimators for modelling eye movements in reading (22). We now turn to the results for each individual type of eye movement (Fig. 2).

**Fig. 1.**
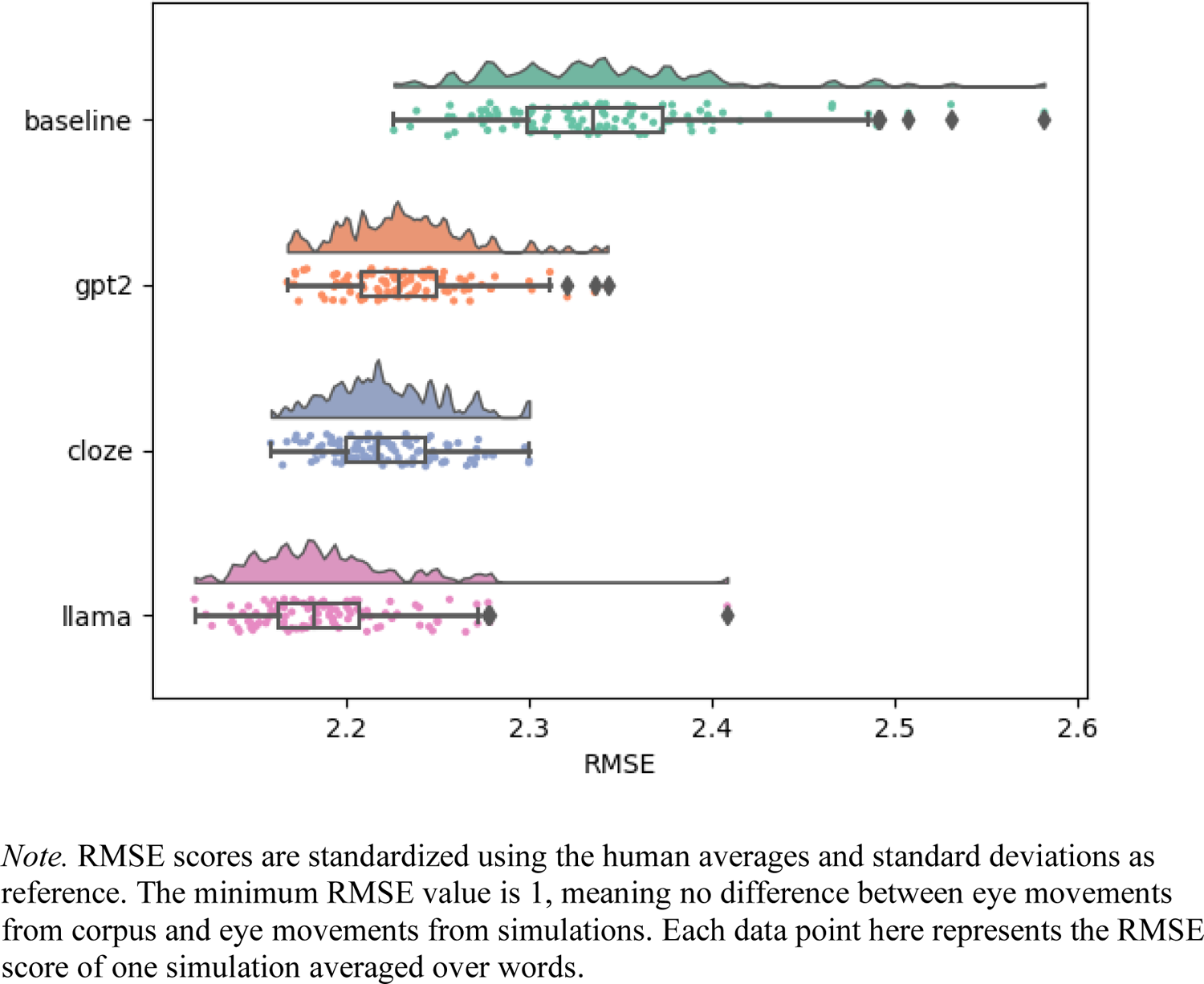
Standardized RMSE scores of OB1 Reader simulations for a baseline without using word predictions, for Cloze-norm predictions and predictions from the GPT2 and LLaMA LLMs.

**Fig. 2.**
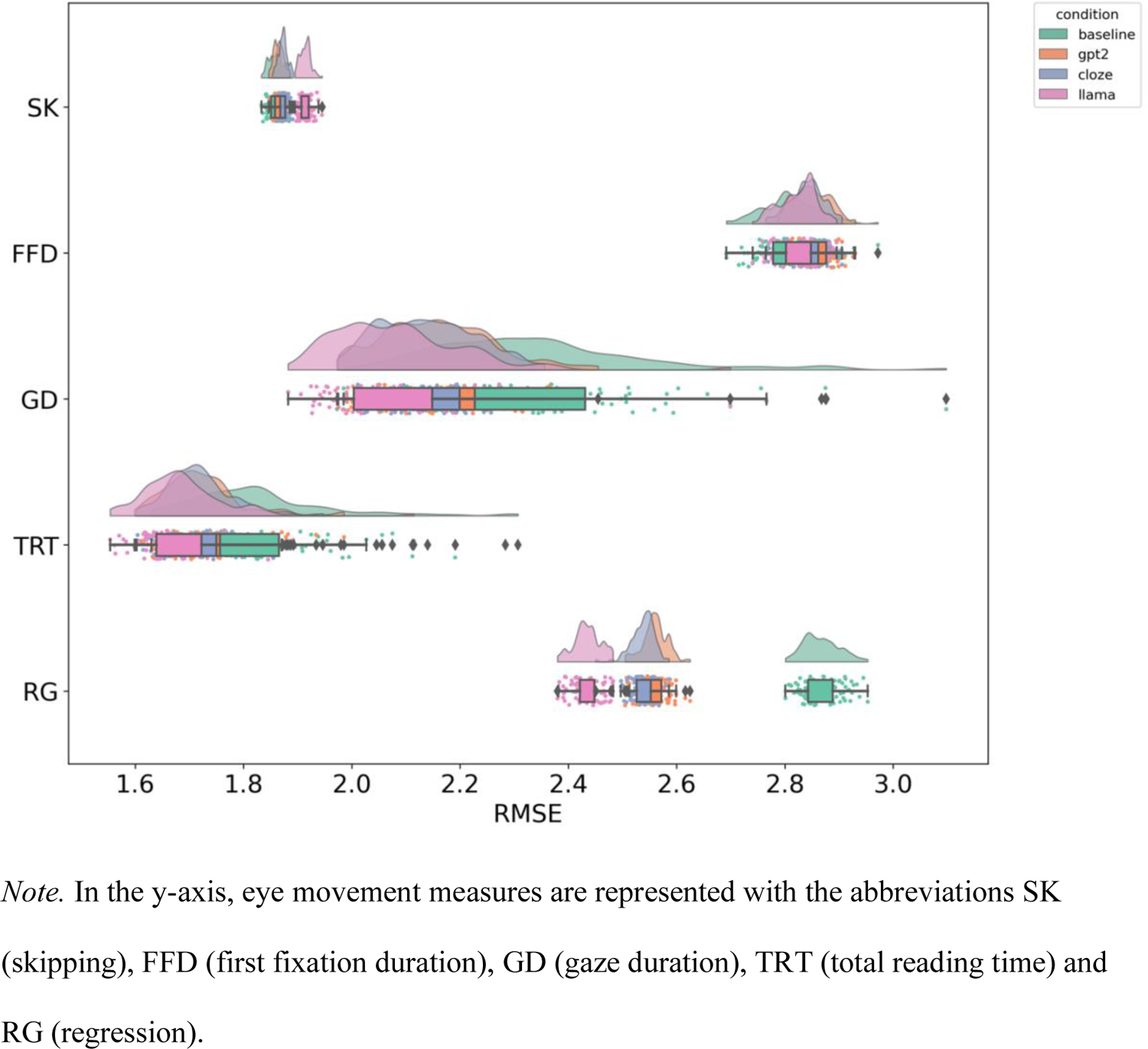
Standardized RMSE scores of OB1-reader simulations per condition for each eye movement measure.

**Fig. 3.**
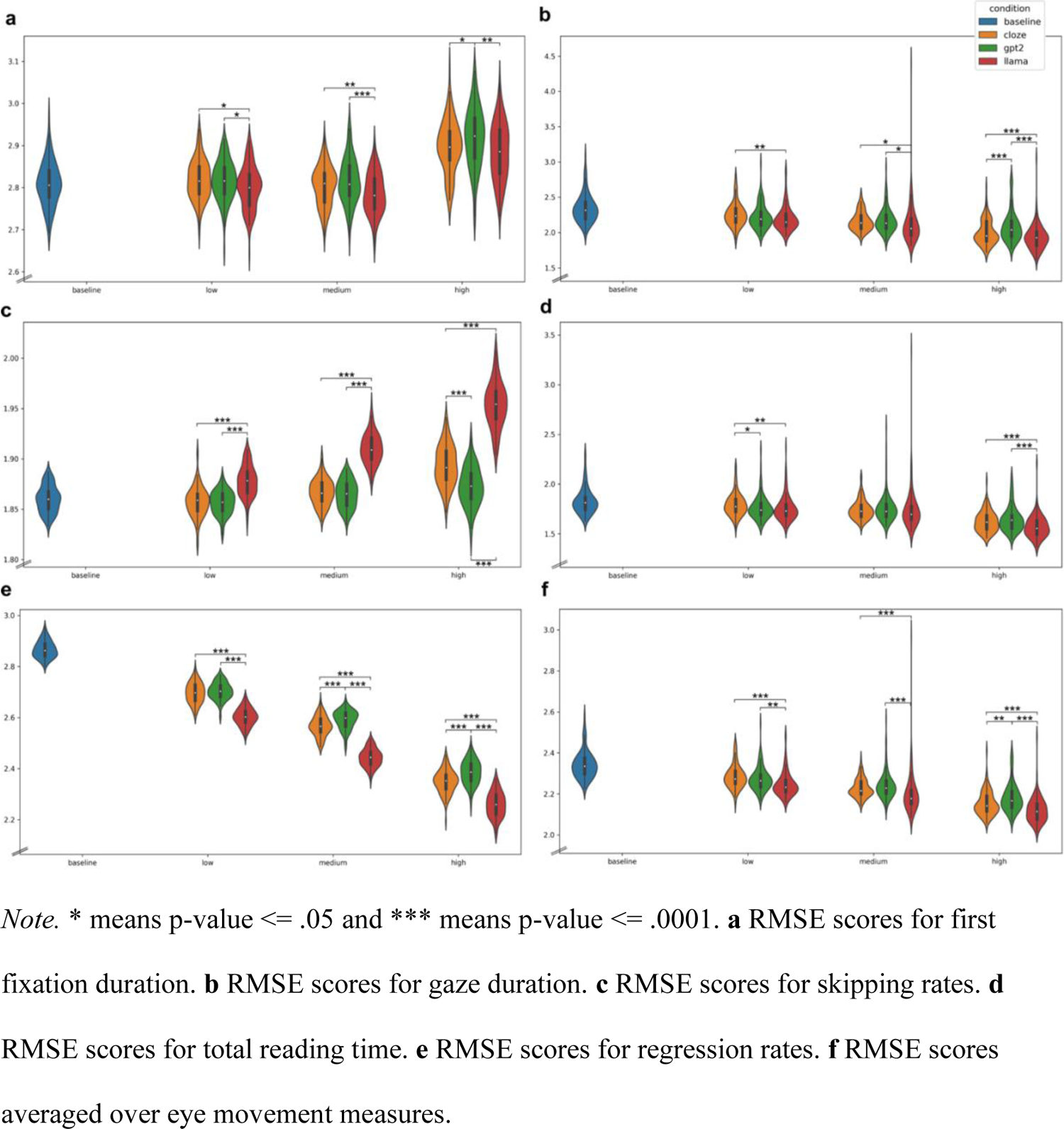
Standardized RMSE scores of OB1-reader simulations per condition, eye movement measure and predictability weight

### First Fixation Duration

RMSE scores for item-level first fixation duration revealed that predictability from LLaMA yielded the best fit compared to GPT2 and Cloze. LLaMA also yielded the least error in each weight condition (see Fig. 3a). When checking predictability effects on first fixation duration (Supporting Information Fig. 1), the relation between predictability and first fixation duration seemed to be weakly facilitatory, with more predictability leading to slightly shorter first fixation duration in both the Provo Corpus and the OB1-reader simulations. This relation occurred in all predictability conditions, suggesting that the LLMs capture a similar relation between predictability and eye movements as Cloze norming, and that this relation also exist for eye movements in the Provo Corpus.

### Gaze Duration

LLaMA produced the least averaged error in fitting gaze duration. GPT2 produced either similar fits to Cloze or a slightly worse fit than Cloze (see Fig. 3b). All predictability conditions reduce error compared to the baseline, confirming the benefit of word predictability on predicting gaze duration. Higher predictability shortened gaze duration (Supporting Information Fig. 2) in both the model simulations (OB1-reader) and in the empirical data (Provo Corpus).

### Skipping

Unexpectedly, skipping rates showed increasing error with predictability compared to the baseline. RMSE scores were higher in the LLaMA condition for all weights (see Fig. 3c). These results show no evidence of skipping being more accurately simulated with any of the predictability estimations tested in this study. While OB1-reader produced sizable predictability effects on skipping rates, these effects seem to be very slight in the empirical data (Supporting Information Fig. 3).

### Total Reading Time

Improvement in RMSE relative to the baseline was seen for total reading time in all conditions. LLaMA showed the best performance, with lower error compared to Cloze and GPT2, especially in the low and high weight conditions (see Fig. 3d). Higher predictability led to shorter total reading time, and this was reproduced by OB1 reader in all conditions. LLaMA showed trend lines for the relation between predictability and total reading time that parallel those seen in the data, suggesting a better qualitative fit than for either Cloze or GPT2 (Supporting Information Fig. 4).

### Regression

Lastly, RMSE for regression was reduced in all predictability conditions compared to the baseline. The lowest error is generated with LLaMA-generated word predictability across predictability weights (see Fig. 3e). Predictability effects on regression are in the expected direction, with higher predictability associated with lower regression rate, but this effect is amplified in the simulations (OB1-reader) relative to the empirical data (Provo Corpus) as indicated by the steeper trend lines in the simulated regression rates (Supporting Information Fig. 5).

## Discussion

The current paper is the first to evidence that large language models (LLMs) can complement cognitive models of reading at the functional level. While previous studies have shown that LLMs provide predictability estimates which can fit reading behaviour as good as- or better than Cloze norming, here we go a step further by showing that language models outperform Cloze norming when used in a cognitive model of reading and tested in terms of simulation fits. Our results suggest that LLMs can provide the basis for a more sophisticated account of semantic processes in models of reading comprehension.

### Word predictability from language models improves fit to eye-movements in reading

Using predictability values generated from LLMs (especially LLaMA) to regulate word activation in a cognitive model of eye-movement control in reading (OB1-reader) reduced error between the simulated eye-movements and the corpus eye-movements, relative to using no predictability or to using Cloze predictability. Late eye-movement measures (total reading time and regression) also showed reduced error in the LLM predictability conditions, with decreasing error the higher the predictability weight. Notably, the benefit of predictability is less clear for early measures (skipping, first fixation duration and gaze duration) than for late measures across conditions. One interpretation of this result is that predictability also reflects the ease of integrating the incoming button-up input with the previously processed context, with highly predictable words being more readily integrated with the previously processed context than highly unpredictable words (23).

Surprisingly, predictability did not improve the fit to skipping. Even though adding predictability increases the average skipping rate (which, importantly, causes the model to better approximate the average skipping rate depicted by humans) there nonetheless appears to be a mismatch between the model and the human data in terms of which individual words are skipped specifically. One potential explanation lies in the effect size of word predictability on skipping, which is higher in the simulations than in the eye-tracking data. Given the model’s assumption that early (i.e. in parafoveal preview) word recognition largely drives skipping behaviour, the boosted effect may have caused highly predictable words to be recognized in the parafovea by the model, while they were not by humans, and highly unpredictable words to not be recognized in the parafovea by the model, while they may have been by humans. It is plausible that lexical retrieval prior to fixation is not the only factor driving skipping behaviour. More investigation is needed into the interplay between top-down feedback processes, such as predictability, and perception processes, such as visual word recognition, and the role of this interaction in saccade programming.

All in all, RMSE between simulated eye movements and corpus eye movements across eye movement measures indicated that LLMs can provide word predictability estimates which are better than Cloze norming at fitting eye movements with a model of reading. Moreover, the least error across eye movement simulations occurred with predictability derived from a larger language model (in this case, LLaMA), relative to a smaller language model (GPT2) and Cloze norming. Previous studies using transformer-based language models have shown mixed evidence for a positive relation between model size and word prediction accuracy and the ability of the predictability estimates to predict human reading behaviour (17,24–26). Our results align with the studies that have found language model quality to positively correlate with the model’s psychometric predictive power (22).

### Language models may aid our understanding of the cognitive mechanisms underlying reading

Improved fits aside, the broader, and perhaps more important, question is whether language models may provide a better account of the higher-order cognition involved in language comprehension (9,19,27,28). Various recent studies have claimed that deep language models offer a suitable “computational framework”, or “deeper explanation”, for investigating the neurobiological mechanisms of language (9,19,20), based on the correlation between model performance and human data. However, correlation between model and neural and behavioural data does not necessarily mean that the model is performing cognition, because the same input-output mappings can be performed by wholly different mechanisms (this is the “*multiple realizability”* principle) (29). Consequently, claiming that our results show that LLMs constitute a “deeper explanation” for predictability in reading would be a logical fallacy. It at best is a failed falsification attempt, which suggests that language models might be useful in the search for explanatory theories about reading.

Caution remains important when drawing parallels between separate implementations, such as between language models and human cognition (30). The question is then how we can best interpret language models for cognitive theory building. If they resemble language processing in the human brain, how so? One option is to frame LLMs as good models of how language works in the brain, which implies that LLMs and language cognition are mechanistically equivalent. This is improbable however, given that LLMs are tools built to perform language tasks efficiently, with no theoretical, empirical or biological considerations about human cognition. It is highly unlikely that language processing in the human brain resembles a Transformer implemented on a serial processor. Indeed, some studies explicitly refrain from making such claims, in spite of referring to language models as a “deeper explanation” or “suitable computational framework” for understanding language cognition (9,19).

Another interpretation is that LLMs resemble the brain by performing the same task, namely to predict the upcoming linguistic input before they are perceived. Prediction as the basic mechanism underlying language is the core idea of Predictive Coding, a prominent theory in psycholinguistics (3) and in cognitive neuroscience (31,32). However, shared tasks do not necessarily imply shared algorithms. For instance, it has been shown that more accuracy on next-word prediction was associated with worse encoding of brain responses, contrary to what the theory of predictive coding would imply (30).

Yet another possibility is that LLMs resemble human language cognition at a more abstract level: both systems encode linguistic features which are acquired through statistical learning on the basis of linguistic data. The similarities are then caused not by the algorithm, but by the features in the input which both systems learn to optimally encode.

Some studies criticize the use of language models to understand human processing altogether. Having found a linear relationship between predictability and reading times instead of a logarithmic relationship, Smith and Levy (8) and Brothers and Kuperberg (33) speculated the discrepancy to be due to the use of n-gram language models instead of Cloze estimations. One argument was that language models and human readers are sensitive to distinct aspects of the previous linguistic context and that the interpretability and limited causal inference of language models are substantial downfalls. However, language models have become more and more powerful in causal inference and still provide a more easily interpretable measure of predictability than does Cloze. Additionally, contextualized word representations show that the previous linguistic context can be better captured by the state-of-the-art language models, compared to simpler architectures such as n-gram models. More importantly, neural networks allow for internal (e.g. architecture, representations) and external (e.g. input and output) probing, which can help adjudicate among mechanistic explanations (20). All in all, language models show good potential to be a valuable tool for investigating higher-level processing in reading. Combining language models, which are built with engineering goals in mind, with models of human cognition, might be a powerful method to test mechanistic accounts of reading comprehension. The current study is the first to apply this methodological strategy.

Finally, we emphasize that the LLM’s success is likely not only a function of the LLM itself, but also of the way in which its outputs are brought to bear in the computational model. The cognitive mechanism that we proposed, in which predictions gradually affect multiple words in parallel, may align better with LLMs than with Cloze norms, because the outputs of the former are based on a continuous consideration of multiple words in parallel, while the outputs of the latter may be driven by a more serial approach.

The optimal method to investigate language comprehension may be by combining the ability of language models to functionally capture higher-order language cognition with the ability of cognitive models to mechanically capture low-order language perception.

Computational models of reading are built as a set of mathematical constructs to define and test explanatory theories or mechanism proposals regarding language processing during reading. As such, they are more interpretable and more resembling of theoretical and neurobiological accounts of cognition than LLM’s. However, they often lack functional generalizability and accuracy. In contrast, large language models are built to efficiently perform natural language processing tasks, with little to no emphasizes on neurocognitive plausibility and interpretability. Interestingly, despite the reliance on performance over explanatory power, LLMs have been shown to capture various aspects of natural language, in particular at levels of cognition considered higher order by brain and language researchers (e.g. semantics and discourse) and which cognitive models of reading often lack. This remarkable ability of LLMs suggests that they offer a promising tool for expanding cognitive models of reading.

## Methods

### Eye-tracking and Cloze Norming

We use the cloze completion and reading time data from the Provo corpus (21). This corpus consists of data from 55 passages (2689 words) with an average of 50 words and 2.5 sentences from various genres, such as online news articles, popular science and fiction. In an online survey, 470 participants provided a cloze completion to each word position in each passage. Each participant was randomly assigned to complete 5 passages, resulting in 15 unique continuations filled in by 40 participants on average. All participants were English native speakers, ages 18-50, with at least some college experience. Another 85 native English-speaking university students read the same 55 passages while their eyes were tracked with a high-resolution, EyeLink 1000 eye-tracker.

The cloze probability of each continuation in the upcoming word position was equivalent to the proportion of participants that provided the continuation in the corresponding word position. Since the number of participants completing a sequence was a maximum of 43, the minimum cloze probability of a continuation was 0.023 (i.e. if each participant would give a different continuation). Words in a passage which did not appear among the responded continuations received cloze probability of 0. The cloze probabilities of each word in each passage and the corresponding continuations were used in the model to pre-activate each predicted word, as further explained on page 23.

The main measure of interest in this study is eye movements. During reading, our eyes make continuous and rapid movements, called *saccades*, separated by pauses in which the eyes remain stationary, called *fixations*. Reading in English consists of fixations of about 250ms on average, whereas a saccade typically lasts 15-40ms. Typically, about 10-15% are saccades to earlier parts of the text, called *regressions*, and about two thirds of the saccades skip words (34).

The time spent reading a word is associated with the ease of recognizing the word and integrating it with the previously read parts of the text(34). The fixation durations and saccade origins and destinations are commonly used to compute word-based measures that reflect how long and how often each word was fixated. Measures that reflect early stages of word processing such as lexical retrieval include first fixation duration (FFD), i.e. the duration of the first fixation on a word, gaze duration (GD), i.e. the sum of fixations on a word before the eyes move forward, and skipping rate (SK). Late measures include total reading time (TRT) and regression rate (RG) and are said to reflect full syntactic and semantic integration(35). Facilitatory effects of word predictability are generally evidenced in both early measures and late measures: that is, predictable words are skipped more often and read more quickly (36).

The measures of interest readily provided in the eye-tracking portion of the corpus were first fixation duration, gaze duration, total reading time, skipping likelihood and regression likelihood. Those measures were reported by the authors to be predictable from the cloze probabilities, attesting the validity of the data collected. Our choice of corpus stemmed from a few advantages of using the Provo corpus in relation to other corpora. Sentences are presented as part of a multi-line passage instead of in isolation (37), which is closer to natural, continuous reading. In addition, Provo provides predictability norms for each word in the text, instead of only the final word (38), which is ideal for studies in which every word is examined. Finally, unlike other cloze corpora that tend to use more constrained contexts which are actually rare in natural reading, this corpus provides more naturalistic cloze probability distributions (21).

### Language Models

Language model probabilities were obtained from two transformer-based language models: the smallest version available of the pre-trained LLaMA (39) (7B parameters, 32 hidden layers, 32 attention heads, 4096 hidden dimensions and 32k vocabulary size); and the smallest version of the pre-trained GPT2 (40) (124M parameters, 12 hidden layers, 12 attention heads, 768 hidden dimensions and 50k vocabulary size). Both models were freely accessible through the Hugging Face Transformers library at the time of the study. The models are auto regressive and thus trained on predicting a word based uniquely on its previous context. Given a sequence as input, the language model computes the likelihood of each word in the model’s vocabulary to follow the input sequence. The likelihood values are expressed in the form of logits in the model’s output vector, where each dimension contains the logit of a corresponding token in the model’s vocabulary. The logits are normalized using softmax operation to be between 0 and 1.

Since the language model outputs a likelihood for each token in the model’s vocabulary, we limited the sample to only the tokens with likelihood above a threshold (0.01). The threshold was defined according to two criteria: the number of predicted words by the language model should be close to the average number of cloze responses over text positions, and the threshold value should be sufficiently low in order to capture the usually lower probabilities of language models.

Each sequence was tokenized with the corresponding model’s tokenizer before given as input, since the language models have their own tokenization (Byte-Pair Encoder (41)) and expect the input to be tokenized accordingly. Pre-processing the tokens and applying the threshold on the predictability values resulted in an average of 10 continuations per word position (range 1 to 26) with LLaMA and an average of 10 continuations per word position (range 1 to 36) with GPT2. After tokenization, no trial (i.e. Provo passage) was longer than the maximum lengths allowed by LLaMA (2048 tokens) nor by GPT2 (1024 tokens). Note that the tokens forming the language model’s vocabulary do not always match a word in OB1-reader’s vocabulary. This is because words can be represented as multi-tokens in the language model vocabulary. Additionally, OB1-reader’s vocabulary is pre-processed and limited to the text words plus the most frequent words in a frequency corpus. 31% of the predicted tokens by LLaMA were not in OB1-reader’s vocabulary and 17% of words in the stimuli are split into multi-tokens in LLaMA’s vocabulary. With GPT2, these percentages were 26% and 16%, respectively.

To minimize the impact of vocabulary misalignment, we considered a match between text word and predicted token when the predicted token corresponded to the first token of the text word as tokenized by the language model tokenizer. For instance, the text word “customary” is split into the tokens “custom” and “ary” by LLaMA. If “custom” is among the top predictions from LLaMA, we used the predictability of “custom” as an estimate for the predictability of “customary”.

### Reading Simulations

In the model of reading simulations OB1-reader (illustrated in Fig. 1), predictive pre-activation exerts a facilitatory effect on word recognition through faster reading times and more likelihood of skipping. Words in the text which are predicted receive additional activation prior to their fixation, which allows for the activation threshold for lexical access to be reached more easily. Consequently, the predicted word may be recognized more readily and even before fixation. In addition, higher predictability may indirectly increase the likelihood of skipping, because more successful recognition in the model leads to a larger attention window. In contrast, pre-activation of predicted words may exert an inhibitory effect on the recognition of words with low predictability, because pre-activated, predicted words inhibit the words that they resemble orthographically, which may include the word that was truly in the text.

**Fig. 1.**
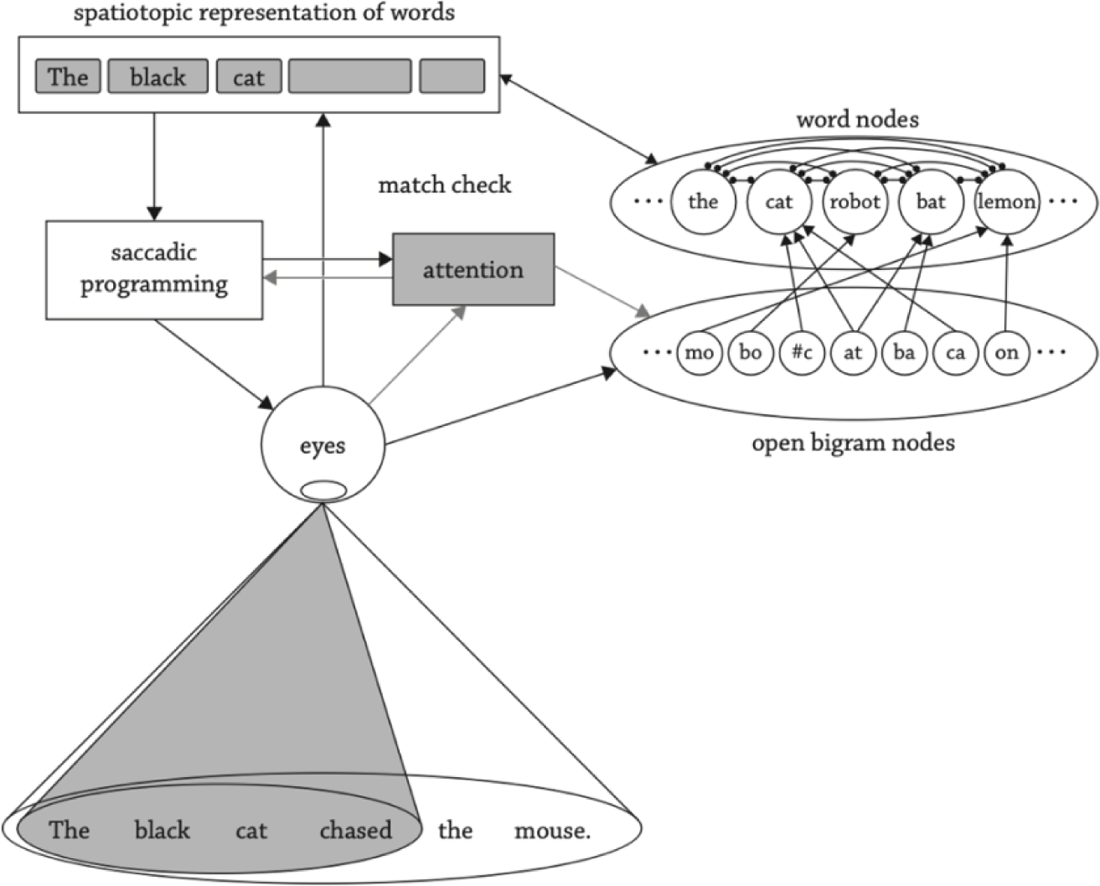
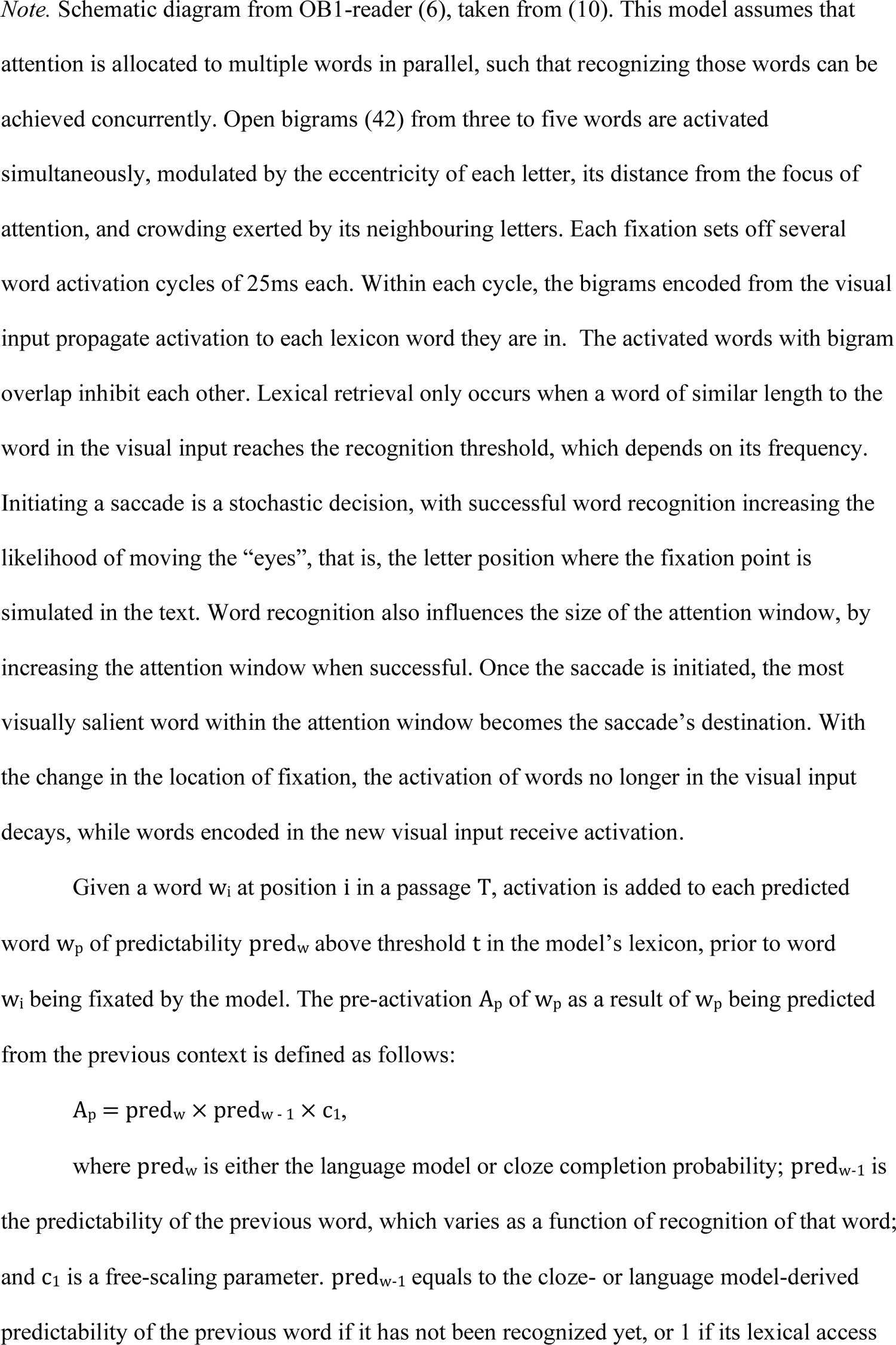
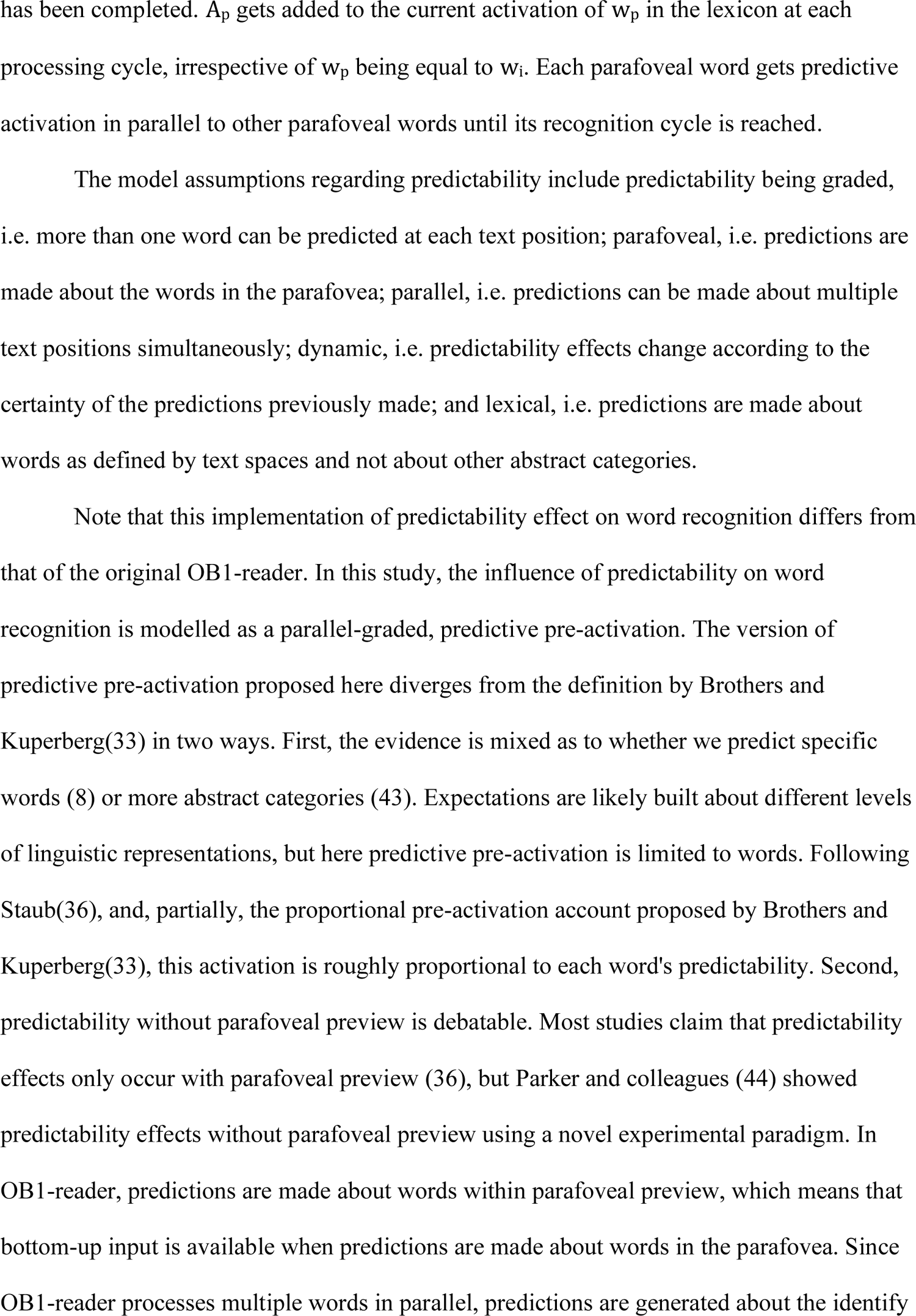
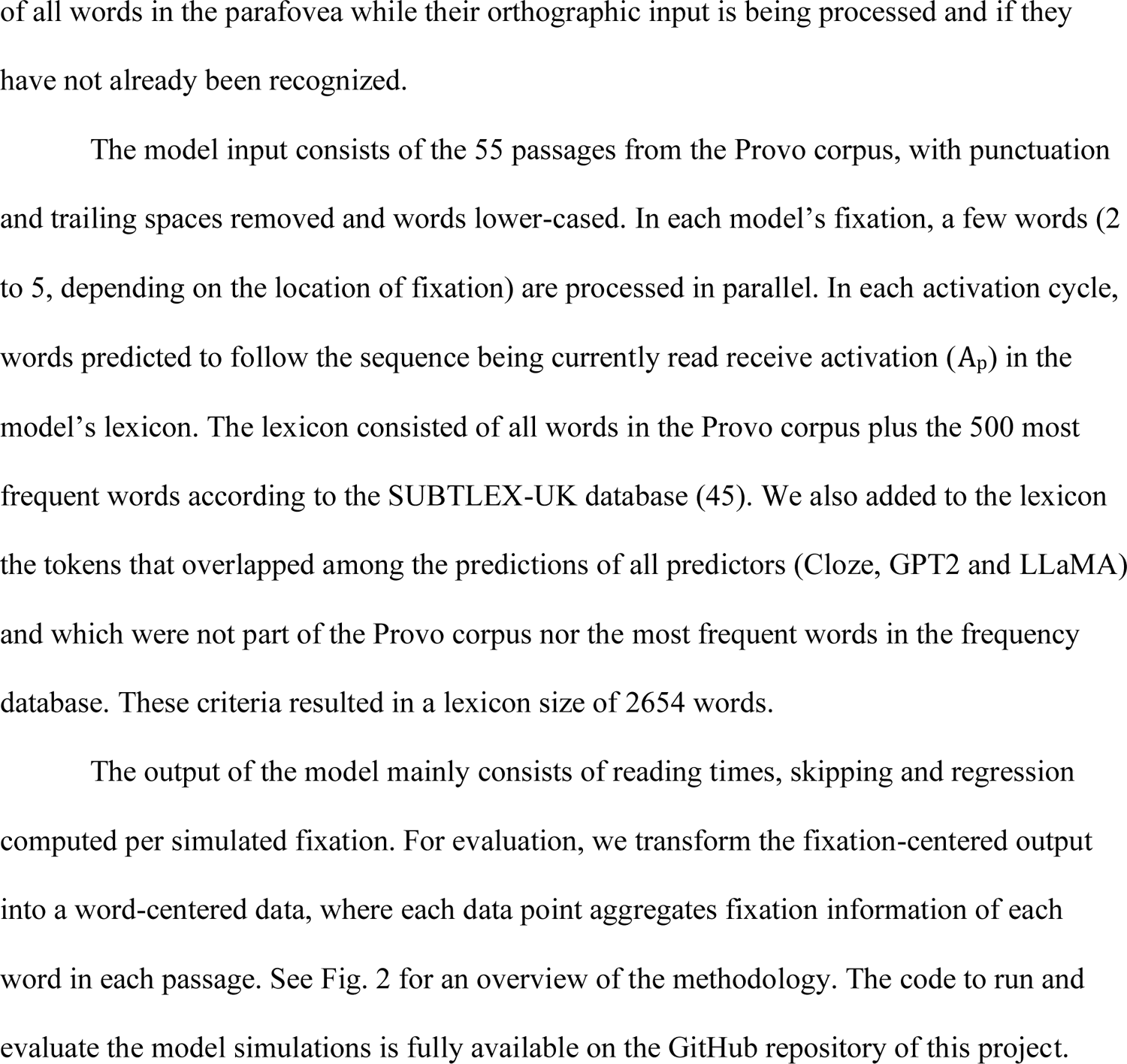

**Fig. 2.**
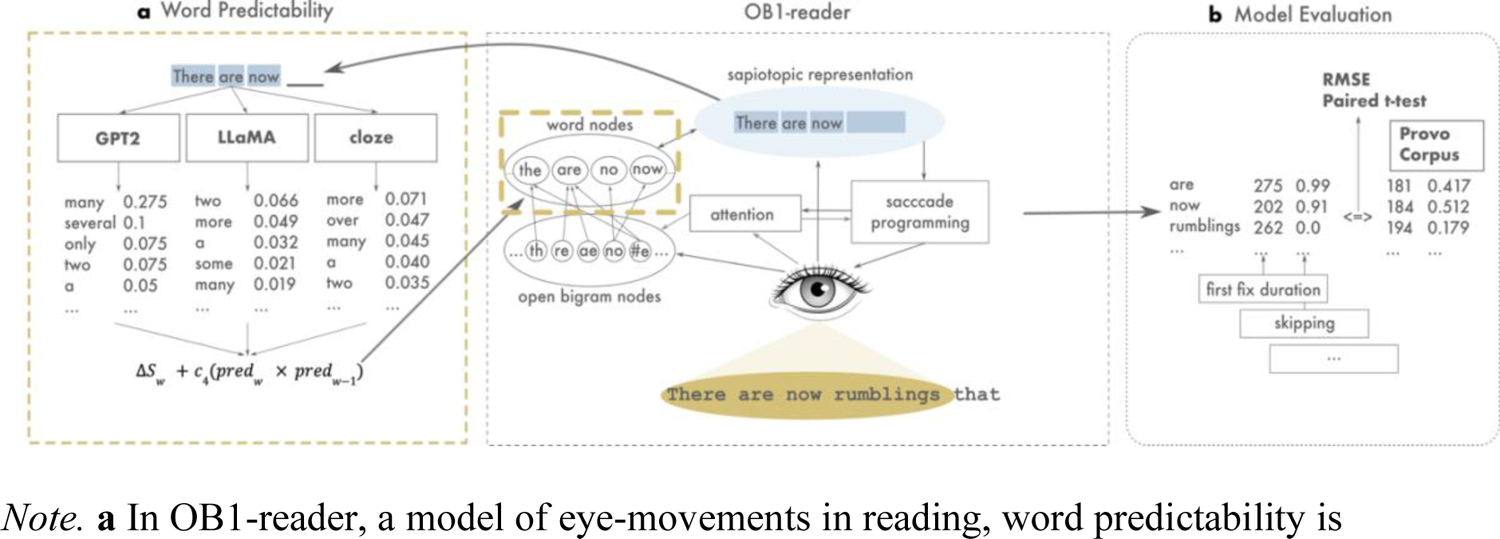
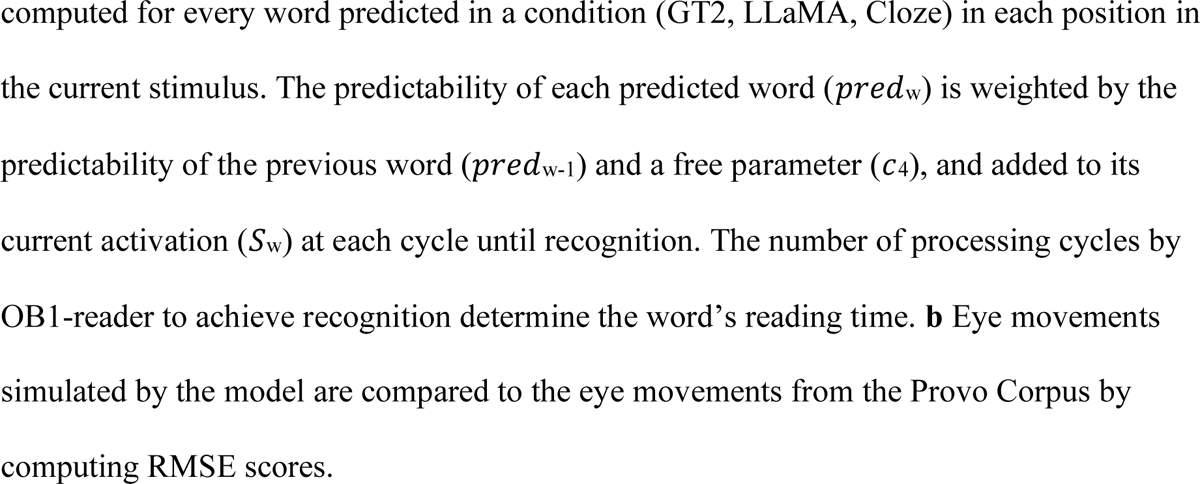
Methodology used in experiment with OB1-reader, LLMs and Cloze Predictability

